# The intestinal microbiota-driven dopamine level influences social interaction and innate color preference in zebrafish larvae

**DOI:** 10.1101/2021.02.26.433134

**Authors:** Ju Wang, Feng Zheng, Lifen Yin, Shengnan Shi, Bing Hu, Lei Zheng

**Affiliations:** Engineering Research Center of Bio-Process, Ministry of Education, School of Food and Biological Engineering, Hefei University of Technology, Hefei, 230009, China; Intelligent Interconnected Systems Laboratory of Anhui Province, Hefei University of Technology, Hefei, 230009, China; School of Life Science, University of Science and Technology of China, Hefei, 230027, China

**Keywords:** Intestinal microbiota, Social interaction, Innate color preference, Dopamine, Zebrafish

## Abstract

Gut microbiota influence neurodevelopment of brain and programing of behaviors. However, the mechanism underlining the relationship between shoals’ behaviors and intestinal microbiota remain controversial and the roles of neurotransmitters are still unclear. Here we show that, shoaling behavior affected the innate color preference of shoals, indicating that shoals tended to choose a favorable color environment that benefits social contact. Meanwhile, administration of D1-R antagonist disrupted the social interaction which led to the deficits of color preference. More importantly, the altered microbiota caused by an antibiotic OTC decreased the sociability and weakened shoals’ color preference. When given a supplement of LGG after OTC exposure, fish exhibited an unexpectedly recovery capability in social cohesion and color preference. Our findings show that dopamine level of brain could mediate both social recognition and color preference, and highlight the pathway of microbial metabolites through the microbiota-gut-brain axis that coordinate the production of dopamine.

## Introduction

Microorganisms, especially the microbiota which reside in the gastrointestinal tract, may influence brain physiology and social behaviors across diverse animal species (**Sherwin et al., 2019**). Social behavior is observed only when animals gather into a group. Some of these behaviors are beneficial for animals, such as division of labor, cooperative care, shared risk and increased immunity, and others are deleterious for groups, such as conflict, dominance and coercion (**Mason and Shan, 2017**). Microbes are implicated in determining social cues in animals, and the altered gut microbiota may affect the social interactions between animals (**Buffington et al., 2016**). For example, germ-free mice which is absence of the microbiota displays deficits in social cognition and social recognition (**Sgritta et al., 2019; Desbonnet et al., 2014**). A successful strategy in unraveling the role of the microbiota in social behavior is the use of antibiotics to deplete the microbiota. A mixture of antibiotics can reduce the social cohesion of zebrafish (**Wang et al., 2016**). Although the perturbations of the microbiota often negatively affect social interaction, modulation of gut microbiota through probiotic supplement can have a beneficial effect. For instance, the social deficit can be reversed during treatment with *Lactobacillus reuteri* (**Buffington et al., 2016**), and zebrafish with the probiotic strain *Lactobacillus rhamnosus* IMC 501 increase shoaling behavior (**Borrelli et al., 2016**). Because social interaction is such a vital component of human and animals’ mental health, the change of this behavior may have negative effects on the whole fitness (**Holt-Lunstad et al., 2015**). Deficits in social interaction manifest in several neuropsychiatric diseases such as schizophrenia, social anxiety, and depression (**Sherwin et al., 2016**).

As is well known, zebrafish are social animals and tend to travel in shoals (**Miller and Gerlai, 2007**). Meanwhile, there is a vital necessity for fish to sense danger and stay away from predators, thus it is beneficial for a fish to stay tight within a shoal and to assess the social interaction efficiently. Furthermore, animals in groups are often better able to perceive colors in their environment (**Endler, 1992; Kelber and Osorio, 2010; Skorupski and Chittka, 2011**), and their color preferences are either most represented in the environment (**Lunau et al., 2011**) or contrast with the background (**Endler et al., 2005**). Plenty of researches have proven that zebrafish have preference towards different colors (**Avdesh et al., 2012; Bault et al., 2015; Oliveira et al., 2015; Fleisch and Neuhauss, 2006; Li et al., 2013; Colwill et al., 2005**). Surprising, the color preference of zebrafish has been extensively studied but still remains controversial. For instance, some researches show a strong preference for blue (**Avdesh et al., 2012; Fleisch and Neuhauss, 2006**), whereas others report a clear aversion for this color (**Bault et al., 2015; Oliveiraet al., 2015; Li et al., 2013; Colwill et al., 2005**). It is still not clear whether the shoaling behavior affect the color preference. We make a hypothesis that the characteristic of background color may influence the efficiency of social contact within a shoal and consequently shoal may prefer certain background colors in order to keep the social contact, which can be called as innate color preference of shoals. Recently, Park *et al*. (**2016**) used zebrafish larvae (5 days post fertilization (dpf)) to test the innate color preference in shoals. However, shoaling behavior usually starts to develop after 7 dpf, becoming progressively stronger for the mature (**Buske and Gerlai, 2011; Dreosti, 2015; Hinz, 2017**). As far as we know, no research has been published that investigate the innate color preference of shoals, although shoaling and social behavior in general has received considerable critical attentions.

The cohesion of shoals has been found to be associated with the whole brain dopamine level (**Buske and Gerlai, 2012**). Dopamine is one of the major neurotransmitters in the central nervous system of the vertebrate brain which plays important roles in a variety of cerebral functions, such as mood, attention, reward, and memory (**Ma and Lopez, 2003; Girault and Greengard, 2004; Vidal-Gadea et al., 2011**). Abundant evidence shows that dopamine is associated with the neurobehavioral functions in zebrafish (**Saif et al., 2013; Shams et al., 2018**). Saif *et al.* (**2013**) found that strong social stimuli will increase the dopamine and its metabolite 3,4-dihydroxyphenylacetic acid (DOPAC) levels in the brain of the adult zebrafish. The short-term isolated zebrafish could reduce the level of DOPAC (**Shams et al., 2018**). The social interaction of shoals in zebrafish can be affected by the dopaminergic system through influence of the dopamine level (**Scerbina et al., 2012**). D1 dopamine receptor antagonist (SCH23390) is most abundantly expressed dopamine receptor subtypes in the brain of zebrafish (**Li et al., 2007**). And SCH23390 disrupts social preference of zebrafish by decreasing the level of dopamine in dopaminergic system (**Steketee, 1998; Kurata and Shibata, 1991**).

In the present study, we evaluated the influence of antibiotic and probiotic which affected gut microbiota of zebrafish on the social interaction and innate color preference. Meanwhile, we assessed the influence of SCH23390 which could affect dopamine level of brains on the social interaction and innate color preference. To analyze the dopamine levels and gut microbial communities, we established the correlation between intestinal microbiota and behaviors through neurotransmitters in brains. Further understanding of how intestinal microbiota influence the brain may be helpful for elucidating the causal mechanisms underlying behavior of shoals and for generating new behavioral paradigms for social disorders, such as social anxiety, and depression.

## Results

### OTC or LGG affects social interaction and color preference in larval zebrafish

In this study, in order to assess the influence of oxytetracycline (OTC) or *Lactobacillus rhamnosus* GG (LGG) on shoal interaction and color preference, the larval zebrafish (5 dpf) were exposed by OTC (500 µg/L) and LGG (10^6^ CFU/mL) for 21 days, respectively (Figure 1A). To investigate the locomotor behavior of larvae which may affect shoaling behavior and color selection, the larvae were transferred into 12-well plates to measure locomotion. the behavioral trajectory and the results of average movement speed (velocity) were not significantly changed by OTC and LGG exposure (Figure 1B-C). The results showed that OTC or LGG may not cause the locomotor change in zebrafish larvae.

**Figure 1.**
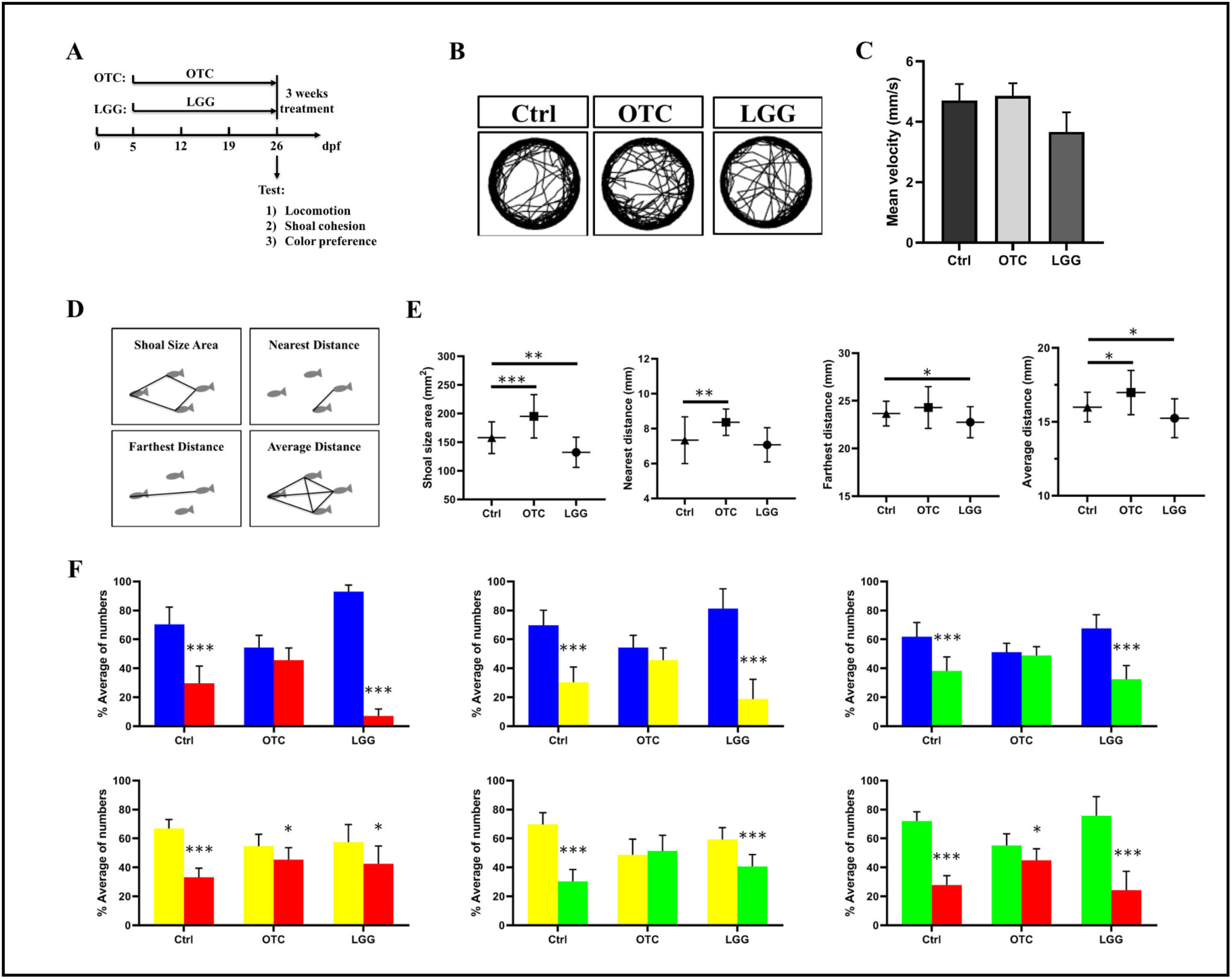
OTC or LGG affects social interaction and color preference in larval zebrafish. (A) Schematic describing the experimental procedure for OTC or LGG treatment. The behavioral trajectory (B) and the mean velocity (C) of the zebrafish larvae from control group (Ctrl) and exposure groups (OTC or LGG) were tested during 10-min period (n=12/ group, 4 groups per treatment). (D) Schematic describing the testing index of social interaction. (F) The shoaling behavior of zebrafish larvae through OTC or LGG treatment (n=6/group, 4 groups per treatment). (G) The shoals of zebrafish larvae exhibit the color preference with 6 color combinations (B-R, B-Y, B-G, Y-R, Y-G, G-R) for OTC or LGG treatment. The percentage of numbers of 20-larval zebrafish spent in each colored zone was counted every 20 s for a total of 10 min (n=10 for Ctrl group, n=8 for OTC group, n=9 for LGG group). * p<0.01, ** p<0.01, *** p<0.001.

The larvae shoaling behavior was analyzed for shoal size area (SA), nearest distance (ND), farthest distance (FD) and average distance (AD) in control (Ctrl), OTC and LGG group (Figure 1D). As demonstrated in Figure 1E, in OTC group, compared to Ctrl group, although FD was not significantly different between the two groups, SA (t_23_ = 4.20, P < 0.001), ND (t_23_ = 3.41, P < 0.01) and AD (t_23_ = 2.78, P < 0.05) significantly increased. Additionally, shoaling behavior of larvae was significantly different between the Ctrl and LGG group. Although ND did not change between the two treatments groups (t_23_ = 0.79, P > 0.05), SA (t_23_ = 3.47, P < 0.01), FD (t_23_ = 2.22, P < 0.05) and AD (t_23_ = 2.30, P < 0.05) were significantly decreased in the LGG group. Together, these data indicated that the shoal cohesion in larvae was decreased through the OTC exposure, and increased through the LGG exposure.

To investigate color preference in shoals with different treatments, a shoal was introduced and allowed to swim freely in color-enriched CPP tank. After 5-min acclimation to the treatments, the location of each larva in each colored zone was counted every 20 s for 10 min total of video recording. As showed in Figure 1F, in Ctrl group, there was a distinct color preference among two color combination (B-R, B-Y, B-G, Y-R, Y-G, G-R), and the order of BYGR preference was B > Y > G > R. However, the OTC treatment group lost their color preference (B-R, B-Y, B-G, Y-G) or attenuated their color selection (Y-R, G-R). Unlike the OTC group, the LGG treatment group maintain the inherent color preference except the preference of Y-R was a little bit decreased, and the order of BYGR preference was the same with Ctrl group (B > Y > G > R). Hence, our results showed that the color preference of shoals was lost or weakened through the OTC treatment, and the LGG exposure group can maintain the existence of the innate color preference in shoals.

### LGG improves the recovery of social interaction and color preference after OTC treatment

The natural microbiota has small amounts of probiotic lactic acid bacteria, and probiotic supplement can reverse the social deficits (**Buffington et al., 2016**). However, whether probiotic can confer neuroprotection or ameliorate deficiency of color preference caused by antibiotic treatment has not been tested. To address this issue, zebrafish larvae received probiotic LGG (10^6^ CFU/ mL) starting immediately after 21 days OTC (500 µg/L) treatment and then again daily until 21 days (Figure 2A).

**Figure 2.**
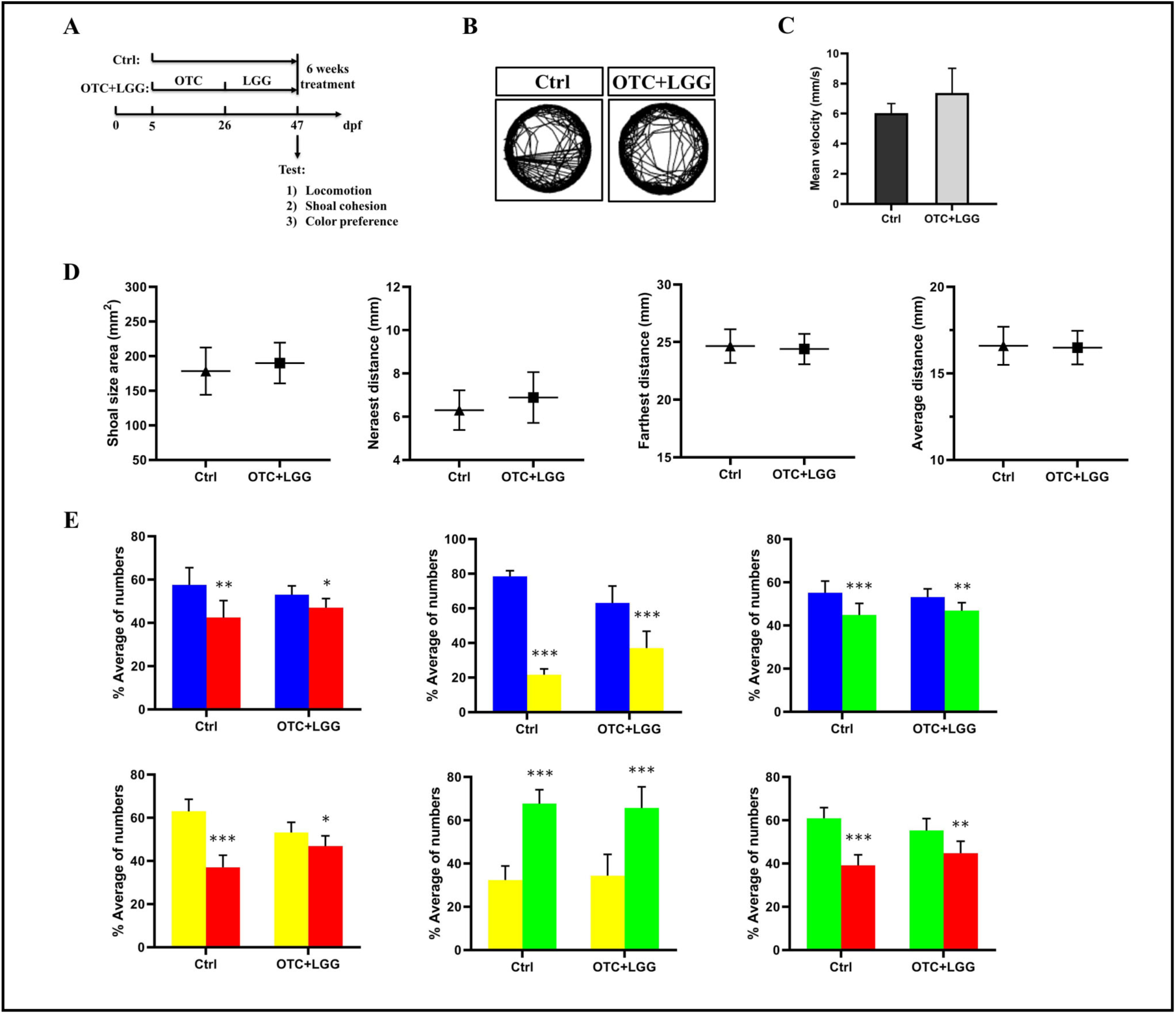
LGG improves the recovery of social interaction and color preference after OTC treatment. (A) Schematic describing the experimental procedure for OTC and LGG treatment. The behavioral trajectory (B) and the mean velocity (C) of the zebrafish larvae from control group (Ctrl) and exposure groups (OTC+LGG) were tested during 10-min period (n=12/ group, 4 groups per treatment). (D) The shoaling behavior of zebrafish larvae through OTC and LGG treatment (n=6/group, 4 groups per treatment). (G) The shoals of zebrafish larvae exhibit the color preference with 6 color combinations (B-R, B-Y, B-G, Y-R, Y-G, G-R) for OTC and LGG treatment. The percentage of numbers of 20-larval zebrafish spent in each colored zone was counted every 20 s for a total of 10 min (n=8 for Ctrl or OTC+LGG group, respectively). * p<0.01, ** p<0.01, *** p<0.001.

From Figure 2B-C, the results showed that the LGG strain could not change the behavioral trajectory and mean velocity after OTC exposure. We next set out to identify the recovery of social cohesion, as showed in Figure 2D, there were no significant difference between Ctrl and OTC+LGG group among shoaling behavior. These data indicated that probiotic LGG could promote the recovery of shoal cohesion through the damage caused by OTC. More importantly, we focused on the variation of color preference, from Figure 2E, although the selections of colors among B-R, B-G, Y-R, and G-R were a little weakened, the order of BYGR preference was the same with Ctrl group (B > G > Y > R). These data raised the possibility that LGG strain could promote the recovery of social cohesion and color selection through the damage caused by OTC.

### SCH23390 affects shoal interaction and color preference in larval zebrafish

From the above results in Figure 1E-F and Figure 2D-E, the strength of social cohesion may drive the color discrimination in shoals of zebrafish. To identify the role of shoaling behavior in color preference, we selected a D1-receptor antagonist SCH23390, which could disrupt social preference and decrease social interaction in zebrafish. To start the experiment, the SCH23390 (SCH) group (a shoal of 20 larvae at 26 dpf) remained in the SCH23390 solution for 1 h (Figure 3A). Because the dopaminergic system is associated with a number of brain functions (e.g., motor function). Thus, D1-receptor antagonist may potentially lead to the altered shoaling behavior as a result of abnormal motor or activity level. However, our results demonstrated that the treat fish did not display abnormal locomotory activity or posture patterns and their mean velocity was also unaltered (Figure 3B-C).

**Figure 3.**
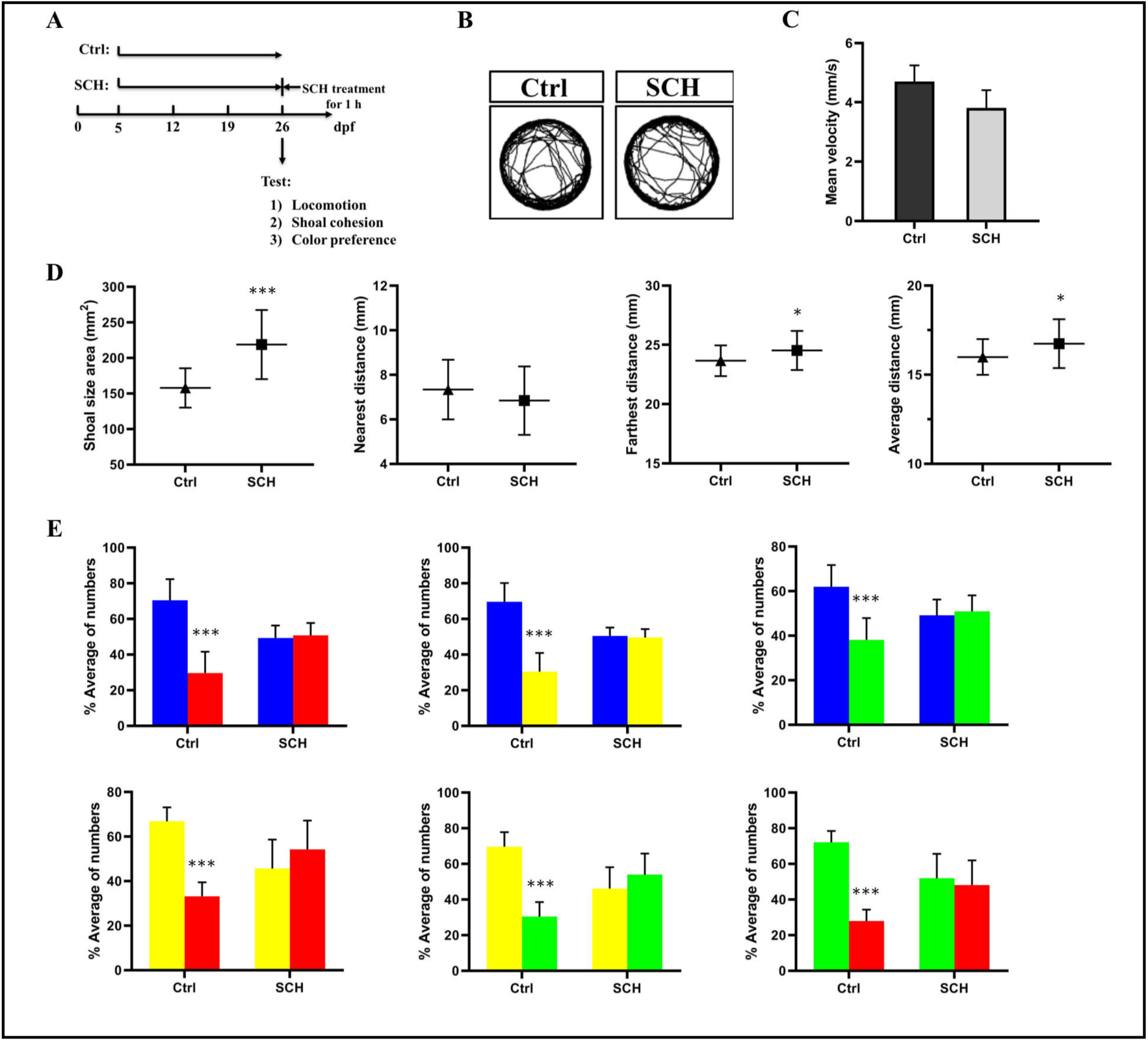
SCH23390 affects shoal interaction and color preference in larval zebrafish. (A) Schematic describing the experimental procedure for SCH23390 treatment. The behavioral trajectory (B) and the mean velocity (C) of the zebrafish larvae from control group (Ctrl) and SCH23390 treatment groups (SCH) were tested during 10-min period (n=12/group, 4 groups per treatment). (D) The shoaling behavior of zebrafish larvae through SCH23390 treatment (n=6/group, 4 groups per treatment). (G) The color preference of shoals with SCH23390 treatment in 6 color combinations (B-R, B-Y, B-G, Y-R, Y-G, G-R). The percentage of numbers of 20-larval zebrafish spent in each colored zone was counted every 20 s for a total of 10 min (n=9 for Ctrl or SCH group, respectively). * p<0.01, ** p<0.01, *** p<0.001.

From the results in Figure 3D, shoal cohesion of larvae treated with SCH23390 was significantly different from Ctrl group. SA (t_23_ = 5.91, P < 0.001), FD (t_23_ = 2.08, P < 0.05), and AD (t_23_ = 2.22, P < 0.05) were significantly decreased in the SCH group, but ND did not alter between the two treatments group (t_23_ = 1.21, P > 0.05). These data suggested that SCH23390 significantly decreased social and explorative behavior in zebrafish larvae. From the results of color preference, by contrast with the shoals without SCH23390 treatment, the shoals lost their interest in color combinations in color-enriched CPP tank (Figure 3E). These results suggested that the color preference of shoals required the participation of social interaction.

### Dopamine level analysis of the treatment groups

As D1-receptor antagonist mediated the social interaction and affected color selection in shoals of larval zebrafish (Figure 3D-E), we hypothesized that the altered social interaction and color preference of the exposed larvae may associated with the dopamine level in the brain. To target the dopamine level, we tested the dopamine concentration of the exposed groups using ELISA. As showed in Figure 4A, OTC was significantly decreased the dopamine level (t_2_ = 6.27, P < 0.05), while LGG was significantly increased the dopamine concentration (t_2_ = 4.82, P < 0.05). From Figure 4B, LGG recover the dopamine level after the OTC treatment (t_2_ = 0.34, P > 0.05). In addition, our data verified the results that SCH23390 can significantly decreased the dopamine level (t_2_ = 37.16, P < 0.001) in the larval brains (Figure 4C). These results suggest that the dopamine which is associated with social cohesion is primary responsible for eliciting the color preference.

**Figure 4.**
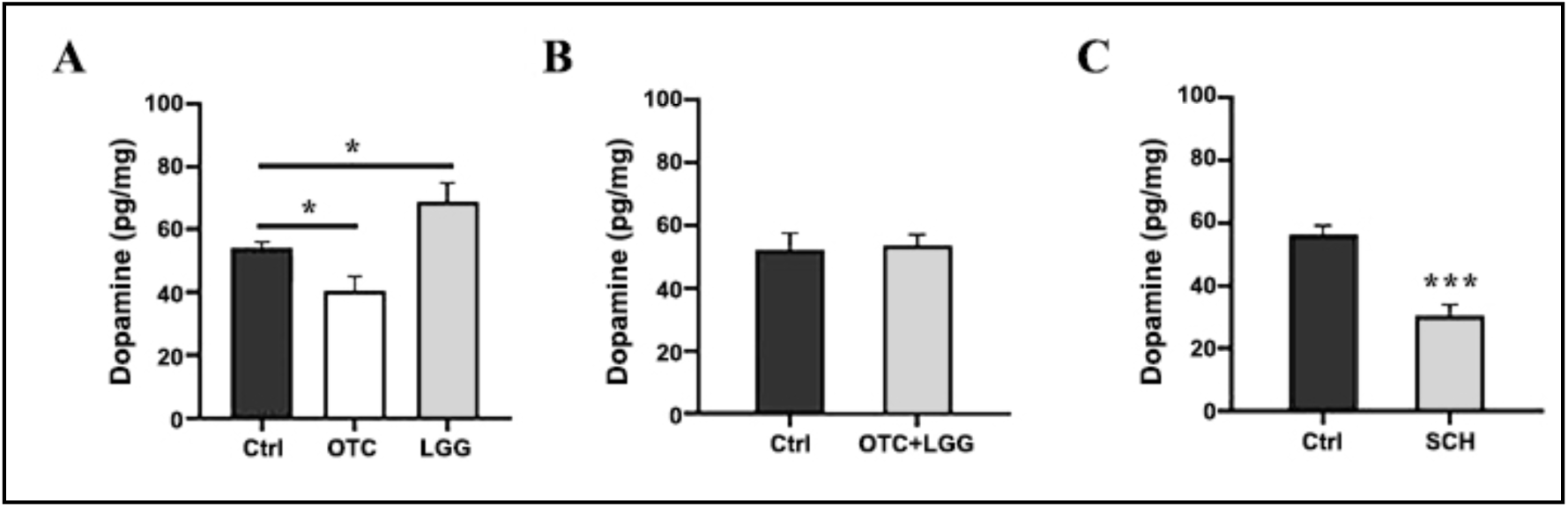
Dopamine level analysis of the treatment groups. (A) The dopamine levels in the brain of larvae through OTC or LGG treatment. (B) The dopamine levels in the brain of larvae through OTC and LGG treatment. (C) The dopamine levels in the brain of larvae through SCH23390 treatment. Error bars represents the standard error of the mean (SEM) (n=30/group, 3 groups). Statistical significance was set to P<0.05 (∗), P<0.01(∗∗), P<0.001 (∗∗∗).

### Gut microbial communities of zebrafish in OTC or LGG treatment group

Considering that the intestine is the main area of antibiotic or probiotic absorption and that intestinal health is very important to fish. The gut microorganisms of the OTC or LGG group were identified using 16S rRNA gene sequencing (Miseq) of the total DNA extracted directly from the two treatment groups. A Venn chart at the OTU level showed the similarity of each group with 152 shared members, and the OTC group showed low similarity compared to the Ctrl (Figure 5A). The community diversity (Shannon) and richness (Chao) were evaluated in Figure 5B-C. Compared with Ctrl group, OTC group had lower bacterial diversity (P < 0.05), and LGG had higher bacterial diversity (P < 0.01) (Figure 5B). The microbial richness from OTC group had a significant decrease compared with the Ctrl group (P < 0.05), and the LGG group showed an increasing trend with the Ctrl group but this was not statistically significant (P > 0.05). PCoA was used to characterize the distribution of variations in community composition among the three treatments, with most of the variation explained by the first two coordinates (22.5% and 36.7%, respectively, Figure 5D). Samples from the OTC and the LGG treatment group were distinctly separated from the Ctrl group, this result indicated that the OTC or LGG had a greater influence on gut bacterial communities.

**Figure 5.**
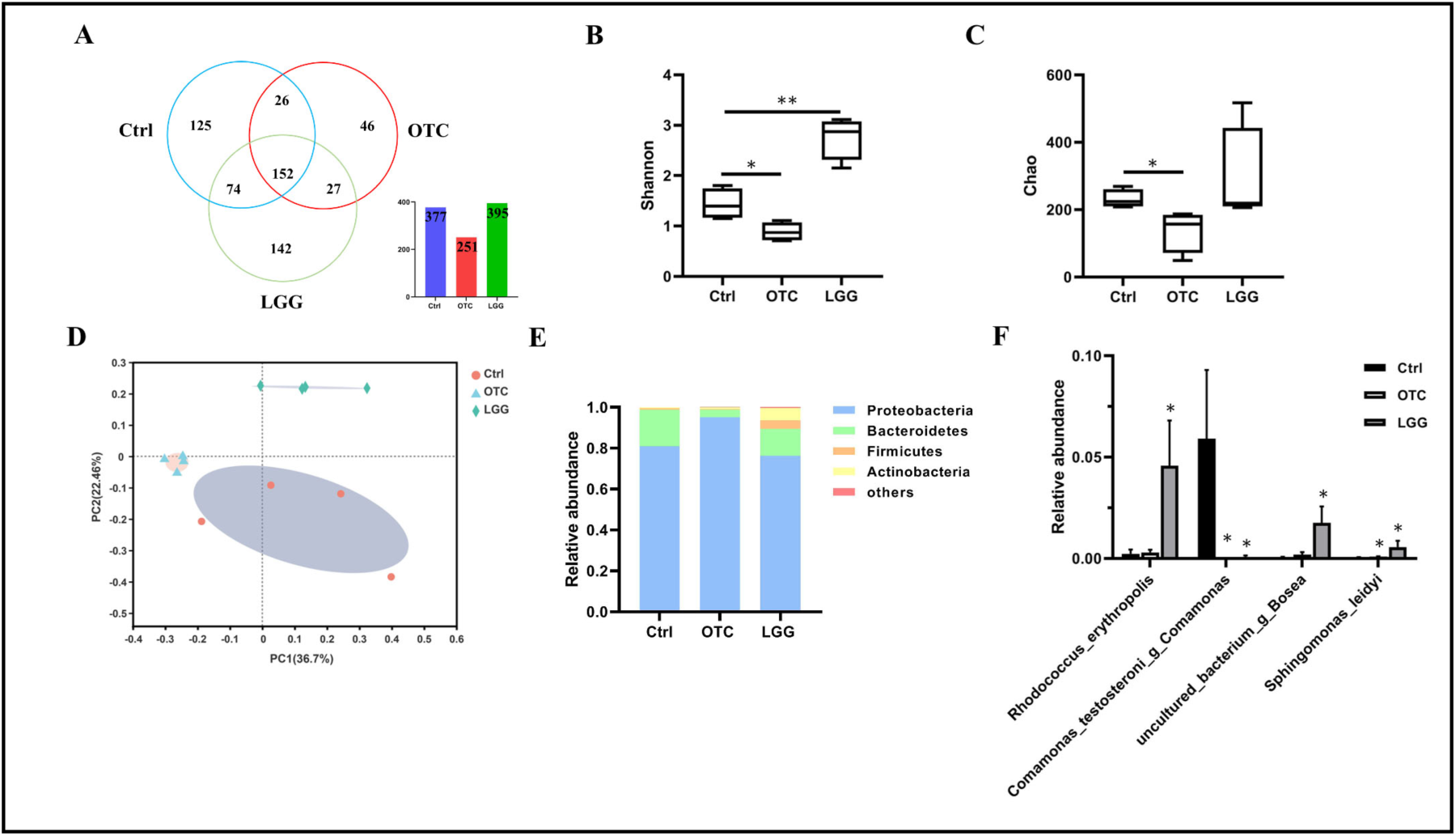
Effect of OTC or LGG exposure on the intestinal microbial composition. (A) The Venn diagram of the three groups. Shannon (B) and Chao (C) index of gut microbiota in the treatment groups. (D) Principal co-ordinates analysis (PCoA) of intestinal bacteria in the treatment groups. Gut microbiota composition at the phylum level (E) and at the species level (F). Error bars represents the standard error of the mean (SEM). Statistical significance was set to P<0.05 (∗), P<0.01(∗∗), P<0.001 (∗∗∗).

The massive sequencing of the gut microbiota revealed that Proteobacteria was the predominant phylum. In Ctrl group, the relative abundances of the four phyla (Proteobacteria, Bacteroidetes, Firmicutes and Actinobacteria) were 81.0%, 17.7%, 0.8%, 0.4%, respectively. In the OTC treatment group, the relative abundances of the four phyla were 95.0%, 3.8%, 0.3% and 0.9%, respectively. In the LGG treatment group, the relative abundances of the four phyla were 76%, 6.4%, 6.6% and 9.8%, respectively. To assess the relative abundance of species level, the relative abundance of the four species is depicted in Figure 5F. *Rhodococcus_erythropolis* was the most abundant bacterium in LGG group, and had a significant increase compared with control group. *Comamonas_testosteroni_g_Comamonas* was found a significant decrease in OTC or LGG group compared with the control. While a significant increase of relative abundance in *uncultured_bacterium_g_Bosea* was observed in LGG group compared to control group. In addition, *Sphingomonas_leidyi* was also found a significant increase in OTC or LGG group compared to the control. Hence, these data indicated that OTC or LGG exposure could affect the intestinal bacterial communities.

### Gut microbial communities of zebrafish in OTC and LGG treatment group

To investigate the role of gut microbiota in promoting the recovery of social cohesion and color selection, we analyzed the composition of zebrafish intestinal microbiota by LGG strain supplementary after OTC treatment. As shown in Figure 6A, the similarity between Ctrl group and OTC+LGG group were 162 shared members in a Venn diagram at the OUT level. From Figure 6B-C, the microbial richness (Chao) and community diversity (Shannon) from the OTC+LGG group showed a decreasing trend with the control group but this was not statistically significant (P > 0.05). Samples from Ctrl and OTC+LGG group clustered together in PCoA (Figure 6D), the data indicated that the OTC+LGG group had a lower influence on gut bacterial communities compared with Ctrl group.

**Figure 6.**
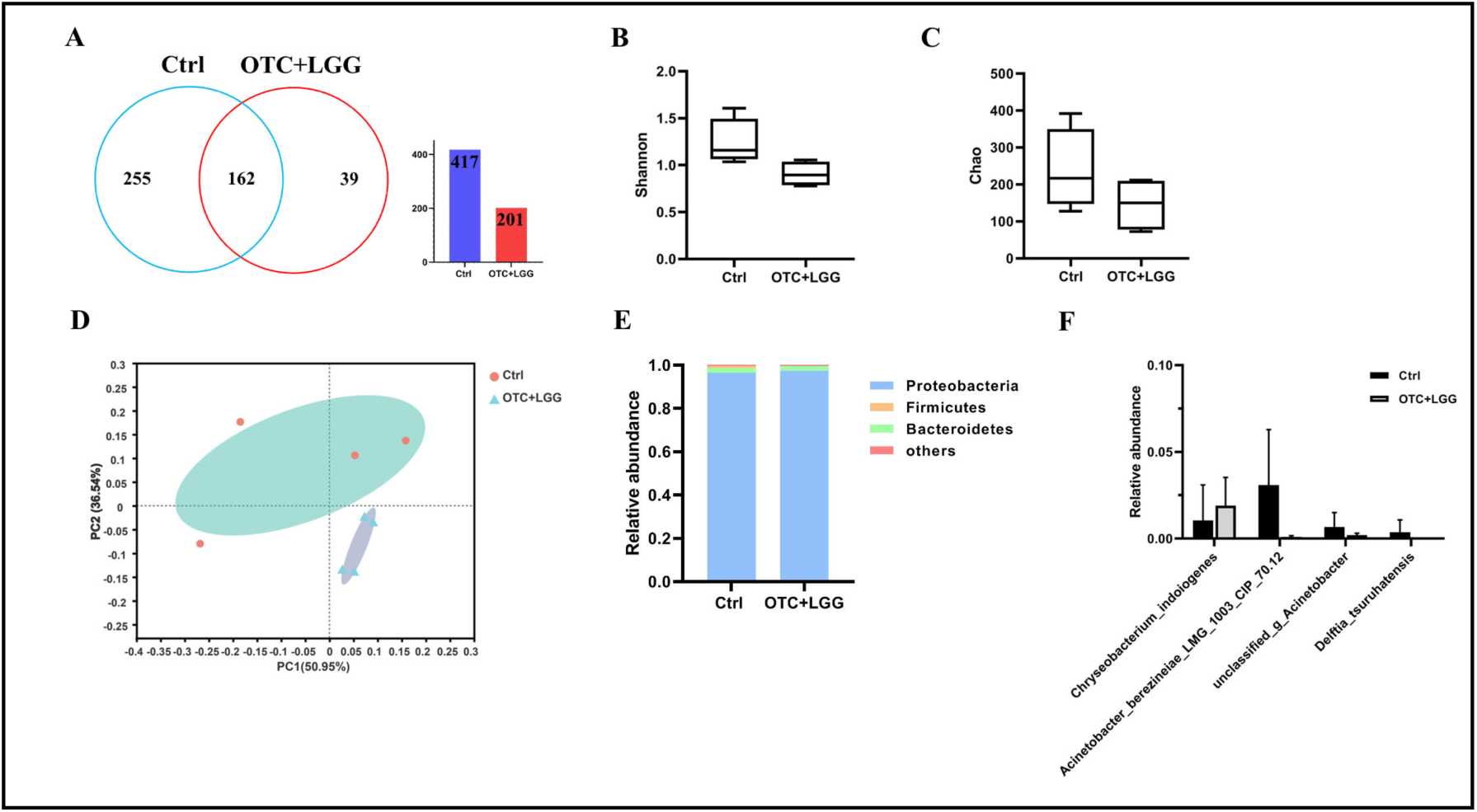
Effect of OTC and LGG exposure on the intestinal microbial composition. (A) The Venn diagram of the two groups. Shannon (B) and Chao (C) index of gut microbiota in the treatment groups. (D) Principal co-ordinates analysis (PCoA) of intestinal bacteria in the treatment groups. Gut microbiota composition at the phylum level (E) and at the species level (F). Error bars represents the standard error of the mean (SEM). Statistical significance was set to P<0.05 (∗), P<0.01(∗∗), P<0.001 (∗∗∗).

The relative abundance of gut microbiota between Ctrl group and OTC+LGG group at phylum level is shown in Figure 6E. The dominant phyla in the two groups were Proteobacteria, Bacteroidetes and Firmicutes. In the control group, the relative abundances of the three phyla were 96.5%, 2.2% and 2.3%, respectively. The relative abundances of the three phyla in OTC+LGG group were 97.4%, 2.1% and < 0.1%, respectively. Additionally, the relative abundance of four species in control and OTC+LGG group was assessed in Figure 6F. The results showed that the relative abundance of four species did not have a significant difference between Ctrl and OTC+LGG group. Therefore, these results suggested that probiotic LGG strain could recovery the gut bacterial composition of zebrafish larvae after the damage by OTC treatment.

## Discussion

The shoaling behavior is one of the most robust and consistent behavioral features in zebrafish, which has been observed both in nature (**Engeszer et al., 2007**) and in the laboratory (**Gerlai, 2014**). Fish often forms aggregations and is found mostly in lakes, puddles, ponds, rice fields, ditches and small watercourses (**Spence et al., 2007**). Sociability varies markedly among different living conditions and associates with gut microbiota. OTC is a commonly used antibiotics in human and veterinary medicine (**Rigos and Troisi, 2005**) which has been frequently detected in surface water, groundwater and seawater (**Nie et al., 2013**). Probiotic as a living and supplementary microorganism, are found in foods (e.g., yogurt, snacks, breakfast cereals and infant formulas) (**Suez et al., 2019**). From our results, the social cohesion of larvae was decreased through OTC treatment, and increased during the LGG treatment (Figure 1E). More importantly, the strain LGG could reversed the social deficits after the treatment with OTC (Figure 2D). Our data of the decreased shoaling behavior were consistent with the study by the antibiotic exposure (β-diketone) (**Wang et al., 2016**). Moreover, studies using probiotics (such as *Lactobacillus reuteri, Lactobacillus rhamnosus* IMC 501) have been beneficial in demonstrating a potential role in regulating social behavior (**Buffington et al., 2016; Borrelli et al., 2016**). Therefore, the changes of living conditions through OTC or LGG exposure may influence sociability and social cognition in zebrafish larvae, and probiotic LGG supplementary could recover the social cohesion after the OTC exposure.

The intricate living conditions which contain abundant colors and shoaling behavior may affect foraging, predator avoidance and reproductive success. The innate color preference would spur shoal on to emigrate to the environment with a favorable color background that benefits social contact. To investigate color preference of zebrafish shoals in different exposure conditions, we measured the color selection of larval shoals with different treatments. According to our studies in Figure 1F, the control group show a clear and strong innate color preference in shoals of larvae, and the order of RYGB preference was B > Y > G > R. In addition, we found that OTC treatment group lost or weaken their color preference. By contrast, the LGG exposure group could maintain the existence of the innate color preference in shoals. To our surprise, probiotic LGG supplementary could recover color selection after the deficiency caused by OTC (Figure 2E). Although the selections of colors among some color combinations were a little weakened, the order of BYGR preference was the same with control group (B > G > Y > R). Some results of color preference were consistent with the findings which were reported by Park *et al*. (**2016**) and Peeters *et al.* (**2016**). They used zebrafish larvae (5 dpf) to test the innate color preference in shoals, and found that zebrafish preferred B over R, and B over Y, and B over G. However, literature has emerged that offers contradictory findings about the innate color preference in shoals. The larvae were employed by different treatments did not exhibit abnormal locomotor behavior or behavioral trajectory (Figure 2B-C, Figure 3B-C). A possible explanation for the contradictory conclusions is that the 5 dpf larvae did not develop shoaling behavior (**Buske and Gerlai, 2011; Dreosti et al., 2015; Hinz and de Polavieja, 2017**). Besides, from our results, the order of color preference in Y-G combination was a little difference between the two control groups at different developmental days. Color is perceived primarily through cones in the retia of zebrafish, and fish do not possess yellow cones which can sense yellow light (**Neitz and Neitz, 2011**), so we hypothesized that the deficits of yellow cones may cause the difference during color selection.

As shown in Figure 1E-F and Figure 2D-E, there was a correlation between social cohesion and color discrimination in shoals of larval zebrafish. To investigate the role of social interaction in the innate color preference in shoals, we employed a SCH23390 and analyzed its effects on behavior of color preference. The drug was chosen because D1-R antagonist, SCH23390, is the most abundantly expressed dopamine receptor subtypes in the brain of zebrafish (**Li et al., 2007**). Second, SCH23390 disrupts social preference of zebrafish by decreasing the level of dopamine in dopaminergic system (**Steketee, 1998; Kurata and Shibata, 1991**). Finally, the drug is water soluble, and zebrafish can be administered by simple immersion in the drug solution. From our results in Figure 3, the D1-R antagonist treatment which can disrupt the social interaction led to the deficits of color preference in shoals. The fish were employed by SCH23390 did not exhibit abnormal motor or posture (Figure 3B-C), and the results of locomotor behavior were consistent with the study by Scerbina *et al.* (**2012**). The dopaminergic system is involved in several brain functions which has been found to be associated with the shoaling tendencies (**Suriyampola et al., 2016**). Dopamine receptors distribute in different brain regions. Clearly, the specific areas are involved in cognition, including hippocampus, the prefrontal cortex, the amygdale, and the ventral and dorsal parts of the striatum. There are four different dopamine receptor subtypes (D1, D2, D3, and D4) in the brain of zebrafish (**Li et al., 2007; Boehmler et al., 2004; Boehmler et al., 2007**). Among the different types of dopaminergic receptors, the excitatory D1 receptor (D1-R) subtype is the most predominately expressed in the brain regions (**Fremeau et al., 1991**). D1-R activate the production of intracellular 3’-5’-cyclic adenosine monophosphate (cAMP) through adenylyl cyclase induction and regulate intercellular calcium signaling or protein kinase activity (**Naderi et al., 2016**). Therefore, these results suggested that social interaction can be affected by dopaminergic system, which could be responsible for eliciting the preference for colors.

We next set out to identify the dopamine level of different treatments that regulate social cohesion and color discrimination in zebrafish larvae. From our results, we found that OTC decreased the dopamine level in brains of larvae, while LGG increased the dopamine concentration (Figure 4A). Meanwhile, LGG recovered the dopamine level after the OTC treatment (Figure 4B). In addition, our data verified the results that SCH23390 severely reduced the dopamine level in the larval brains (Figure 4C). These data raised the possibility that OTC or LGG through influence of dopamine level of larval brains was primary responsible for social cohesion and color preference. The dopamine plays key roles in the neurobehavioral functions in zebrafish (**Saif et al., 2013; Shams et al., 2018**). For instance, the strong social stimuli will increase the dopamine and DOPAC levels in the brain of the adult zebrafish (**Saif et al., 2013**), and the short-term isolated zebrafish could reduce the level of DOPAC (**Shams et al., 2018**). Several potential mechanisms may be responsible for the decreased the level of dopamine through exposure to the SCH23390. The decreased of dopamine levels imply the reduced dopamine production and/or increased dopamine degradation in response to the employed SCH23390 (**Diop et al., 1988**). Yung *et al.* (**1995**) have shown that D1-R localized in the post-synaptic terminals of neurons in the basal ganglia. The blockade of post-synaptic neurotransmitter receptors may impair signaling downstream and reduce neurotransmitter release. The antagonist of D1 receptor could increase the concentration of dopamine in the synaptic cleft which lead to reuptake and leakage to extra-synaptic areas. The increased extra-synaptic dopamine could activate dopaminergic autoreceptors on the pre-synaptic neuron and inhibit the dopamine synthesis (**Tissari and Lillgals, 1993**). Taken together, these studies and our own suggest the role of dopamine level in social interaction and innate color preference of shoals, with the deficits of sociability and color discrimination linked to the decreased dopamine level in the brain of zebrafish.

The altered microbiota, specifically the changes of microbiota that resides in the gastrointestinal system, may influence neurodevelopment and programming of social behaviors (**Sherwin et al., 2019**). Indeed, the antibiotic OTC treatment was associated with a reduction in gut microbiota diversity (Shannon) and richness (Chao) (Figure 5B-C). Moreover, under conditions of OTC exposures, the relative abundance of microbiota on phylum level or in species level was significantly influenced by antibiotic (Figure 5E-F). By contrast, probiotic LGG could promote the increase of gut microbiota diversity and richness (Figure 5B-C), and the relative abundance of microbiota on phylum level (Bacteroidetes, Firmicutes, Actinobacteria) or on species level was increased by LGG treatment (Figure 5E-F). Although antibiotic administration often perturbs microbiota, modulation of gut bacteria through probiotic strain LGG supplementary can have an advantageous result (Figure 6). The intestinal microbiota can signal to the brain which may affect behavioral processes of sociability via numerous pathways (the microbiota-gut-brain axis), including production of microbial metabolites, immune activation, activation of the vagus nerve, and production of various neurotransmitters (**Sherwin et al., 2019**). Gut microbiota produced vast metabolites through the pathway of microbial metabolites, such as volatile carboxylic acids, esters, neurotransmitters (e.g., dopamine), and fatty acids, some of which may influence brain physiology and behavior (**Roshchina, 2016**). Additionally, in vitro studies indicated that certain bacteria had the capacity to produce neurotransmitters such as noradrenaline, dopamine, and - aminobutyric acid (**Taj and Jamil, 2018; Marques et al., 2016**). Some researches indicated that the gut microbiota could influence serotonergic neurotransmission by regulating the availability of its precursor, tryptophan (**Clarke et al., 2013**). Thus, these data raised the possibility that microbiota-driven neurotransmitters (dopamine) mediated both social recognition and color discrimination, with the pathway of microbial metabolites having an important role in the production of dopamine.

## Materials and methods

### Zebrafish husbandry and embryo collection

Zebrafish (AB strain) used in this study were obtained from breeding center at University of Science and Technology of China. Zebrafish were maintained at 28 ± 0.5 ℃ with a 14 h light/10 h dark cycle (room fluorescent light, 08:00 am-22:00 pm). The pH and conductivity in circulating water of the aquarium were 7.0-7.4 and 1500-1600 µs/cm, respectively. All adult zebrafish were fed fresh brine shrimps twice a day, at 09:00 am and 14:00 pm. Prior to the experiment, the embryos were collected from spawning healthy adults. To obtain the normal fertilized embryos for tests, the dead embryos and sundries were cleared.

### Antibiotic and SCH23390 exposure, probiotic culture and exposure

The OTC as a common antibiotic, was purchased from Aladdin (Shanghai, China). OTC was dissolved in water with a concentration at 100 mg/L as a stock solution. Based on the available literature on the concentration of antibiotic which can affect shoaling behavior, the concentration of OTC used in the present study was set at 500 µg/L (**Wang et al., 2016**). The D1-receptor antagonist SCH23390 (R-(+)-8-chloro-2,3,4,5-tetrahydro-3-methyl-5-phenyl-1H-3-benzazepine-1-ol; Cat # D054; Sigma-Aldrich) was applied to treat individuals which significantly reduced the amount of dopamine in the brain of zebrafish (**Scerbina et al., 2012**). Before experiment, fish were placed in drug exposure beaker (0.1 mg/L, 500 mL in volume) and remained in D1-receptor antagonist solution for 1 h. Because the 1 h exposure period is sufficiently long for the drug to reach the zebrafish brain through the vasculature of their gills and skin (**Scerbina et al., 2012**). All the animals in the beaker were offered the same conditions (including illumination, temperature and dissolved oxygen) which were identical to the standard aquarium.

The probiotic strain LGG was obtained from Bnbio (Beijing, China), and was cultured in MRS medium at 37 ℃ for 24 h. The fermentation broth was centrifuged at 1000 rpm for 2 min at 4 ℃. The strain was obtained and washed twice with sterile PBS buffer and resuspended. The final concentration of LGG was 10^8^ CFU/mL and stored at 4 ℃. In order to maintain constant OTC or LGG concentration for the period of exposure, the OTC or LGG solutions were renewed with freshly prepared solutions every day.

### Locomotor activity test

Locomotor behavior was assessed according to the published protocol (**Zhou et al., 2014**). The larval zebrafish were transferred into 12-well plates with one larva and 2 mL system water for each well. The individuals were acclimated to the recording condition for 5 min. After the acclimation, locomotor behavior was tested in a 10-min period. The trajectory of the larva was recorded by industrial camera (Guangdong, China) and the videos were analyzed using LSRtrack and LSRanalyze software which was described by Zhou *et al.* (**2014**). The velocity was recorded to assess motor function.

### Shoaling behavioral assay

To test the social behavior, a four-fish assay was used to measure the social interaction of zebrafish larvae in a 6-well plate according to Kyzer *et al*. (**2012**). Briefly, 24 normal larvae were randomly selected from each treatment and transferred to the 6-well plate with each well containing 4 larvae and 5 mL water. After the acclimation of 5 min, the shoaling behavior was video-recorded for 10 min, and analyzed using 10 screenshots made every 1 min during the recording period. Each screenshot was calibrated to the size of the 6-well plate and measured the distances between each fish in the group using Image J software. The shoal size area, nearest distance, farthest distance, and average distance were calculated to assess the shoaling behavior (**Shen et al., 2020; Borrelli et al., 2016**).

### Color preference test

The color-enriched conditional place preference (CPP) apparatus, is a customized fish tank (20 cm length × 10 cm width × 10 cm height), colored with four color combinations (red (R), green (G), yellow (Y) and blue (B)). To create the preference for two colors, the CPP tank was divided into two compartments which were covered with the corresponding colors on all side except the top. A video camera (NVH-589MW; Wang Shi Wu You Corporation; China) was placed above the CPP tank for vertical video tracking. The color preference apparatus was placed over the LED light panel to ensure the light source could cover the whole tank. The detailed apparatus has been described in our previous work (**Wang et al., 2014**). To test the color preference of shoals in zebrafish larvae, the experimental tanks were poured into 500 mL fresh fish water. There was no physical barrier between the two compartments and the 20-larval zebrafish (a shoal) could swim freely in the entire tank. After 5-min adaption, the proportion of numbers stayed in each colored zone was recorded every 20 s for 10-min experiment. The outside of color-enriched CPP tank was opaque to prevent external visual interference from all direction. To minimize the effect of noise, the experimental room was closed and kept quiet, and the experimenter was not visible to the fish during the recording.

### Dopamine content analysis

Fish were decapitated and the brains of each treatment groups were dissected on ice, and frozen in a microcentrifuge tube (30 brain per tube) at −80 ℃. The level of dopamine was measured by ELISA according to the manufacturer’s (Wuhan Xinqidi Biological Technology Co. Ltd., China) instructions. Briefly, the pre-cooled phosphate buffered saline (PBS, 1:9, m/v) was added to the brain tissues of larvae, the tissues were ground and centrifuged for 2 min at 5000 g to obtain the supernatant. The standards and the supernatant of each sample were added to each well of 48-well plates precoated with primary antibodies. The plates were incubated at 37 ℃ for 60 min after adding enzymeconjugate, and then was rinsed with distilled water. During 30 min of the chromogenic reaction, the Optical Density (O.D.) was measured at 450 nm by a microtiter plate reader (Thermo, USA). The concentration of dopamine (pg/mg brain tissue) is calculated by comparing the O.D. of the samples to the standard curve.

### Bacterial genomic DNA extraction

In order to study the bacterial composition analysis, total bacterial DNA was isolated from 30 larvae per treatment group by using the E.Z.N.A. ^®^ Soil DNA Kit (OMEGA, USA). DNA yield was measured in a NaNo DROP 2000 Spectrophotometer (Thermo, USA).

### Illumina high-throughput sequencing of barcoded 16S rRNA genes

The V3-V4 region of the 16S rRNA gene was amplified with the primers 338F (5’-ACTCCTACGGGAGGCAGCAG-3’) and 806R (5’-GGACTACHVGGGTWTCTAAT-3’). PCR reactions were performed in a reaction mixture containing (20 µL), 4 µL of 5 × Fast Pfu Buffer, 2 µL of 2.5 mM dNTP, 0.8 µL of 5 µM Forward Primer, 0.8 µL of 5 µM Reverse Primer, 0.4 µL of FastPfu Polymerase and 10 ng of template DNA. The PCR consisted of an initial denaturation time of 3 min at 95 ℃ followed by 29 cycles (95 ℃ for 30 s, 55 ℃ for 30 s and 72 ℃ for 45 s), and the final extension period lasted for 10 min at 72 ℃. PCR products were purified and were subjected to Illumina based high-throughput sequencing (Majorbio Bio-Pharm Technology, Co., Ltd., Shanghai, China).

### Bioinformatics analysis

Alpha diversity was calculated by using Phylogenetic Distance over OTU counts (**Faith, 1992**), and the beta diversity was estimated by using Unifrac metric (**Lozupone et al., 2006**). Taxonomic richness and diversity estimators were determined based on OTU abundance matrices in Mothur (Version v.1.30.1), Chao was used to assess the community richness, and the diversity was reflected by Shannon. Principal co-ordinates analysis (PCoA) was the distance matrix and performed by using a MATLAB R2016a environment. The data in taxonomy of intestinal bacterial communities were presented as mean ± standard error of the mean (SEM). Statistical analysis was performed by the two-tailed t test.

### Statistical analysis

All experimental results were expressed as the means ± standard error of the mean (SEM) and analyzed by an independent t-test using the SPSS statistics program. Significance was set at p < 0.05 for all the experiments.

## Acknowledgements

This study is supported by the National Key Research and Development Plan of China (2017YFC1600603), and Funds for Huangshan Professorship of Hefei University of Technology (407-037019).

## Notes

### Competing Interest Statement

The authors have declared no competing interest.

## References

Avdesh A, Martin-Iverson MT, Mondal A, Chen M, Askraba S, Morgan N, Lardelli M, Groth DM, Verdile G, Martins RN. 2012. Evaluation of color preference in zebrafish for learning and memory. Journal of Alzheimer’s Disease. 28: 459–469.

Bault ZA, Peterson SM, Freeman JL. 2015. Directional and color preference in adult zebrafish: Implications in behavioral and learning assays in neurotoxicity studies. Journal of Applied Toxicology 35: 1502–1510.

Boehmler W, Carr T, Thisse C, Thisse B, Canfield VA, Levenson R. 2007. D4 Dopamine receptor genes of zebrafish and effects of the antipsychotic clozapine on larval swimming behavior. Genes Brain & Behavior 6: 155–166.

Boehmler W, Obrecht-Pflumio S, Canfield V, Thisse C, Thisse B, Levenson R. 2004. Evolution and expression of D2 and D3 dopamine receptor genes in zebrafish. Developmental Dynamics 230: 481–493.

Borrelli L, Aceto S, Agnisola C, De Paolo S, Dipineto L, Stiling RM, Dinan TG, Cryan JF, Menna LF, Fioretti A. 2016. Probiotic modulation of the microbiota-gut-brain axis and behavior in zebrafish. Scientific Reports 6: 30046.

Buffington SA, Di Prisco GV, Auchtung TA, Ajami NJ, Petrosino JF, Costa-Mattiolo M. 2016. Microbial reconstitution reverses maternal diet-induced social and synaptic deficits in offspring. Cell 165: 1762–1775.

Buske C, Gerlai R. 2011. Shoaling develops with age in Zebrafish (*Danio rerio*). Progress in Neuropsychopharmacology Biological Psychiatry 35: 1409–1415.

Buske C, Gerlai R. 2012. Maturation of shoaling behavior is accompanied by changes in the dopaminergic and serotoninergic systems in zebrafish. Developmental Psychobiology 54: 28–35.

Clarke G, Grenham S, Scully P, Fitzgerald P, Moloney RD, Shanahan F, Dinan TG, Cryan JF. 2013. The microbiome-gut-brain axis during early life regulates the hippocampal serotonergic system in a sex-dependent manner. Molecular Psychiatry 18: 666–673.

Colwill RM, Raymond MP, Ferreira L, Escudero H. 2005. Visual discrimination learning in zebrafsh (*Danio rerio*). Behavioural Processes 70: 19–31.

Desbonnet L, Clarke G, Shanahan F, Dinan TG, Cryan JF. 2014. Microbiota is essential for social development in the mouse. Molecular Psychiatry 19: 146–148.

Diop L, Gottberg E, Brière R, Grondin L, Reader TA. 1988. Distribution of dopamine D1 receptors in rat cortical areas, neostriatum, olfactory bulb and hippocampus in relation to endogenous dopamine contents. Synapse 2: 395–405.

Dreosti E, Lopes G, Kampff AR, Wilson SW. 2015. Development of social behavior in young zebrafish. Frontiers in Neural Circuits 9: 39.

Endler JA. 1992. Signals, signal conditions, and the direction of evolution. American Naturalist 139: S125–S153.

Endler JA, Westcott DA, Madden JR, Robson T. 2005. Animal visual systems and the evolution of color patterns: sensory processing illuminates signal evolution. Evolution 59: 1795–1818.

Engeszer RE, Patterson LB, Rao AA, Parichy DM. 2007. Zebrafish in the wild: a review of natural history and new notes from the field. Zebrafish 4: 21–40.

Faith DP. 1992. Conservation evaluation and phylogenetic diversity. Biological Conservation 61: 1–10.

Fleisch VC, Neuhauss SC. 2006. Visual behavior in zebrafish. Zebrafish 3: 191–201.

Fremeau RT, Jr Duncan GE, Fornaretto MG, Dearry A, Gingrich JA, Breese GR, Caron MG. 1991. Localization of D1 dopamine receptor mRNA in brain supports a role in cognitive, affective, and neuroendocrine aspects of dopaminergic neurotransmission. Proceedings of the National Academy of Sciences of the United States of America 88: 3772–3776.

Gerlai R. 2014. Social behavior of zebrafish: From synthetic images to biological mechanisms of shoaling. Journal of Neuroscience Methods 234: 59–65.

Girault JA, Greengard P. 2004. The neurobiology of dopamine signaling. Archives of Neurology 61: 641–644.

Hinz RC, de Polavieja GG. 2017. Ontogeny of collective behavior reveals a simple attraction rule. Proceedings of the National Academy of Sciences of the United States of America 114: 2295–2300.

Holt-Lunstad J, Smith TB, Baker M, Harris T, Stephenson D. 2015. Loneliness and social isolation as risk factors for mortality: a meta-analytic review. Perspectives on Psychological Science A Journal of the Association for Psychological Science 10: 227–237.

Kelber A, Osorio D. 2010. From spectral information to animal colour vision: Experiments and concepts. Proceedings Biological Sciences 277: 1617–1625.

Kurata K, Shibata R. 1991. Effects of D1 and D2 antagonists on the transient increase of dopamine release by dopamine agonists by means of brain dialysis. Neuroscience Letters 133: 77–80.

Kyzar EJ, Collins C, Gaikward S, Green J, Roth A. 2012. Effects of hallucinogenic agents mescaline and phencyclidine on zebrafish behavior and physiology. Progress in Neuro-psychopharmacology and Biological Psychiatry 37: 194–202.

Li P, Shah S, Huang L, Carr AL, Gao Y, Thisse C, Thisse B, Li L. 2007. Cloning and spatial and temporal expression of the zebrafish dopamine D1 receptor. Developmental Dynamics 236: 1339–1346.

Li X, Liu B, Li XL, Li YX, Sun MZ, Chen DY, Zhao X, Feng XZ. 2013. SiO2 nanoparticles change colour preference and cause Parkinson’s-like behaviour in zebrafish. Scientific Reports 4: 1–9.

Lozupone C, Hamady M, Knight R. 2006. UniFranc-an online tool for comparing microbial community diversity in a phylogenetic context. BMC Bioinformatics 7: 371.

Lunau K, Papiorek S, Eltz T, Sazima M. 2011. Avoidance of achromatic colours by bees provides a private niche for hummingbirds. Journal of Experimental Biology 214: 1607–1612.

Ma PM, Lopez M. 2003. Consistency in the number of dopaminergic paraventricular organ-accompanying neurons in the posterior tuberculum of the zebrafish brain. Brain Research 967: 267–272.

Marques TM, Patterson E, Wall R, Sullivan OO, Fitzgerald GF, Cotter PD, Dinan TG, Cryan JF, Ross RP, Stanton C. 2016. Influence of GABA and GABA-producing lactobacillus brevis DPC 6108 on the development of diabetes in a streptozotocin rat model. Beneficial Microbes 7: 409–420.

Mason P, Shan H. 2017. A valence-free definition of sociality as any violation of inter-individual independence. Proceedings of the Royal Society B Biological Sciences 284: 20170948.

Miller N, Gerlai R. 2007. Quantification of shoaling behaviour in zebrafish (*Danio rerio*). Behavioural Brain Research 184: 157–166.

Naderi M, Jamwal A, P.Chivers D, Niyogi S. 2016. Modulatory effects of dopamine receptors on associative learning performance in zebrafish (*Danio rerio*). Behavioural Brain Research 303: 109–119.

Neitz J, Neitz M. 2011. The genetics of normal and defective color vision. Vision Research 51: 633–651.

Nie XP, Liu BY, Yu HJ, Liu WQ, Yang YF. 2013. Toxic effects of erythromycin, ciprofloxacin and sulfamethoxazole exposure to the antioxidant system in Pseudokirchneriella subcapitata. Environmental Pollution 172: 23–32.

Oliveira J, Silveira M, Chacon D, Luchiari A. 2015. The zebrafish world of colors and shapes: Preference and discrimination. Zebrafish 12: 166–173.

Park JS, Ryu JH, Choi TI, Bae YK, Lee S, Kang HJ, Kim CH. 2016. Innate color preference of zebrafish and its use in behavioral analyses. Moleculer Cells 39: 750–755.

Peeters BWMM, Moeskops M, Veenvliet ARJ. 2016. Color preference in *Danio rerio*: effects of age and anxiolytic treatments. Zebrafish 13: 330–334.

Rigos G, Troisi GM. 2005. Antibacterial agents in Mediterranean finfish farming: a synopsis of drug pharmacokinetics in important euryhaline fish species and possible environmental implications. Reviews in Fish Biology & Fisheries 15: 53–73.

Roshchina VV. 2016. New trends and perspectives in the evolution of neurotransmitters in microbial, plant, and animal cells. Advances in Experimental Medicine & Biology 874: 25–77.

Saif M, Chatterjee D, Buske C, Gerlai R. 2013. Sight of conspecific images induces changes in neurochemistry in zebrafish. Behavioural Brain Research 243: 294–299.

Scerbina T, Chatterjee D, Gerlai R. 2012. Dopamine receptor antagonism disrupts social preference in zebrafish: A strain comparison study. Amino Acids 43: 2059–2072.

Sgritta M, Dooling SW, Buffington SA, Momin EN, Francis MB, Britton RA, Costa-Mattioli M. 2019. Mechanisms underlying microbial-mediated changes in social behavior in mouse models of autism spectrum disorder. Neuron 101: 246–259.

Shams S, Amlani S, Buske C, Chatterjee D, Gerlai R. 2018. Developmental social isolation affects adult behavior, social interaction, and dopamine metabolite levels in zebrafish. Developmental Psychobiology 60: 43–56.

Shen Q, Truong L, Simonich MT, Huang C, Tanguay RL, Dong Q. 2020. Rapid well-plate assays for motor and social behaviors in larval zebrafish. Behavioural Brain Research 391: 112625.

Sherwin E, Bordenstein SR, Quinn JL, Dinan TG, Cryan JF. 2019. Microbiota and the social brain. Science 366: eaar2016.

Sherwin E, Sandhu KV, Dinan TG, Cryan JF. 2016. May the force be with you: the light and dark sides of the microbiota-gut-brain axis in neuropsychiatry. CNS Drugs 30: 1019–1041.

Skorupski P, Chittka L. 2011. Is colour cognitive? Optics & Laser Technology 43: 251–260.

Spence R, Fatema M K, Ellis S, Ahmed ZF, Smith C. 2007. Diet, growth and recruitment of wild zebrafish in Bangladesh. Journal of Fish Biology 71: 304–309.

Steketee JD. 1998. Injection of SCH 23390 into the ventral tegmental area blocks the development of neurochemical but not behavioral sensitization to cocaine. Behavioural Pharmacology 9: 69–76.

Suez J, Zmora N, Segal E, Elinav E. 2019. The pros, cons, and many unknowns of probiotics. Nature Medicine 25: 716–729.

Suriyampola PS, Shelton DS, Shukla R, Roy T, Bhat A, Martins EP. 2016. Zebrafish social behavior in the wild. Zebrafish 13: 1–8.

Taj A, Jamil N. 2018. Bioconversion of tyrosine and tryptophan derived biogenic amines by neuropathogenic bacteria. Biomolecules 8: 10.

Tissari AH, Lillgäls MS. 1993. Reduction of dopamine synthesis inhibition by dopamine autoreceptor activation in striatal synaptosomes with in vivo reserpine administration. Journal of Neurochemistry 61: 231–238.

Vidal-Gadea A, Topper S, Young L, Crisp A, Kressin L, Elbel E, Maples T, Brauner M, Erbguth K, Axelrod A, Gottschalk A, Siegel D, Pierce-Shimomura JT. 2011. Caenorhabditis elegans selects distinct crawling and swimming gaits via dopamine and serotonin. Proceedings of the National Academy of Sciences of the United States of America 108: 17504–17509.

Wang J, Liu CH, Ma F, Chen W, Liu J, Hu B, Zheng L. 2014. Circadian clock mediates light/dark preference in zebrafish (*Danio Rerio*). Zebrafish 11: 115–121.

Wang X, Zheng Y, Zhang Y, Li J, Zhang H, Wang H. 2016. Effect of β– diketone antibiotic mixtures on behavior of zebrafish (*Danio rerio*). Chemosphere 144.

Yung KKL, Bolam JP, Smith AD, Hersch SM, Ciliax BJ, Levey AI. 1995. Immunocytochemical localization of D1 and D2 dopamine receptors in the basal ganglia of the rat: Light and electron microscopy. Neuroscience 65: 709–730.

Zhou YZ, Cattley RT, Cario CL, Bai Q, Burton EA. 2014. Quantification of larval zebrafish motor function in multiwall plates using open-source MATLAB applications. Nature Protocols 9: 1533–1548.

